# A systematic review of scientific research focused on farmers in agricultural adaptation to climate change (2008-2017)

**DOI:** 10.1101/2020.01.24.917864

**Authors:** Yao Yang, Barnabas C. Seyler, Miao Feng, Ya Tang

## Abstract

Due to the severe consequences of climate change, associated risks to global food security, and the contribution of agriculture to greenhouse gas emissions, agriculture must necessarily adapt to meet these challenges. Many studies have therefore sought to investigate agricultural adaptation to climate change, and as key stakeholders in agriculture, farmers play a vital role in this process. There is a rapidly increasing corpus of scholarship on agricultural adaptation to climate change, with many studies beginning to incorporate survey methods to examine farmer perceptions and adaptation responses. Nevertheless, in-depth understanding of farmers worldwide is inadequate due to insufficiently robust methodologies, socio-economic disparities, and unequal geographic distribution. In this study, we searched and reviewed the existing peer-reviewed, English-language scientific articles published between 2008 and 2017 on agricultural adaptation to climate change that have incorporated farmers into their research methodologies. The main findings include the following: (1) a small but increasing number of studies focus on farmers in climate change adaptation; (2) the global geographic distribution of the reviewed studies is uneven, and many of the most vulnerable nations (e.g., lower-income/agricultural-dependent economies) have no representation at all; (3) there were diverse rationales and methods for incorporating farmers into the studies, and many of the methodological differences were due to practical and logistical limitations in lower-income/agricultural-dependent nations; and (4) studies were from multiple academic fields, indicating the need for more interdisciplinary collaboration moving forward because agricultural adaptation to climate change is too complex for a single discipline to fully explore. Although English is increasingly recognized as the “international language of science,” due to the challenge of language segmentation limiting broader understanding of global scholarship whenever possible, future reviews should be jointly conducted in both English and non-English languages.

## Introduction

Agriculture and climate change are interrelated [1, 2]. On the one hand, agriculture is a ‘victim’ of climate change, since it is sensitive to differences in climatic conditions, so climate change inevitably has significant effects on agriculture [3, 4]. Specifically, climate- and weather-related disruptions can cause extraordinary impacts on food production and price, potentially exacerbating food insecurity for millions globally [5, 6]. Widely cited estimates indicate that from 1980 to 2008, the global yields of two major food crops, namely, wheat and maize, dropped by 5.5% and 3.8%, respectively, due to the impact of climate change [7]. On the other hand, agriculture is also a substantial ‘contributor’ to climate change because it is a significant source of greenhouse gases (GHGs) [8–10]. It is estimated that agriculture generates over 20% of total global anthropogenic GHG emissions [3, 11].

Because the impacts of climate change on agriculture represent threats to quality of life on both the local and global scales [4], and because agriculture has the potential to reduce GHGs by advancing carbon storage technologies [12, 13], wise land use management [14], and well-managed cropping and grazing systems/practices [8, 15–18], calls for the development and implementation of integrated climate change response strategies/policies for agriculture are increasingly urgent [19–21]. As farmers are the key stakeholders in agriculture, the impacts of climate change on agriculture have posed a direct threat to their livelihoods and well-being worldwide [22, 23]. Therefore, a greater understanding of farmers’ beliefs [17, 24] and perceptions [25, 26] of climate change; their concerns about existing/potential climate change impacts [17, 27]; and their preferences [28], willingness [29, 30] and supportiveness [31] of climate change adaptation/mitigation is valuable to both scholars and policy-/decision-makers. In particular, understanding farmers’ beliefs and perceptions of these matters is necessary to properly develop and effectively implement climate change response strategies/policies for the agricultural sector [20, 21, 32].

Scientific publications on the effects of climate change have increased rapidly in recent decades [33]. The Intergovernmental Panel on Climate Change (IPCC) also produces a comprehensive assessment of the existing body of climate change research outcomes on an approximately five-year basis. In addition, the two most renowned international scientific journals, *Nature* and *Science,* publish regular updates on climate change research [34–39]. Many other studies on various topics have focused on agriculture and climate change adaptation [33, 40, 41]. However, the majority of existing studies on agricultural adaptation to climate change are focused on the biophysical and socio-economic perspectives [42–46], and many of these are concentrated on weather responses [47–49], agricultural vulnerabilities [50–54], or agricultural resilience [55–58].

Because of the importance of farmers [59], we expected that many studies would incorporate farmers in their research methodologies and address one or more related concerns in their research objectives. Indeed, our initial literature review indicated that there were thousands of studies related to ‘farmers’ and ‘climate change adaptation,’ and numerous studies have incorporated survey methods to examine farmers’ perceptions and attitudes towards adaptation [41]. However, since a significant number of these studies were conducted or dominated by scholars from natural sciences backgrounds, the incorporation of social science methods into these studies tended to be cursory add-ons to methodologies that are otherwise largely used in the natural sciences. Certainly, all of these studies are valuable and important additions to the existing body of knowledge about agriculture and climate change. However, to increase the utility of these studies and to draw more generalizable conclusions, studies should strive to more equitably incorporate perspectives from both the social sciences and the natural sciences into their methodologies [41]. As such, current scholarship has a considerable weakness in how it incorporates the major stakeholders of agriculture and climate change, namely, farmers, into their studies, particularly from the social science perspective.

To better understand the current status of this weakness, we designed a study to systemically review the published scientific literature related to agriculture and climate change adaptation to assess how many studies and to what extent they have incorporated farmers into their research methodologies. The following questions guided our study: What is the extent of current scholarship that explores farmers’ roles in agricultural adaptation to climate change? By what methods are farmers incorporated into these studies? What are the interests and objectives of climate change studies that do investigate farmers? Where in the world do the scholars and farmers being studied come from? Do the origins of scholars and farmers indicate a potential geographic or socio-economic bias in current scholarship that should be addressed?

## Materials and Methods

Bibliometric studies that analyse trends in research through published literature have been increasingly recognized methods identifying both the focal points of research and their developments as valuable and important [60]. Many reviews have summarized the scientific knowledge of agriculture and climate change and have assessed how agricultural systems might or could adapt to climate change. To gain a clearer picture of the broader body of scholarship related to farmers and climate change adaptation, we conducted a systematic literature review [61]. A systematic literature review involves reviewing articles according to clearly formulated questions, using intentional and explicit criteria to guide the selection of relevant studies and to conduct a comprehensive analysis [62, 63]. A key strength of this approach is its ability to produce a rapid, comprehensive, and transparent knowledge assessment of a specific topic associated with climate change and agricultural adaptation research.

### Literature selection

All papers in this review were collected from the Web of Science Core Collection platform from Thomson Reuters. Web of Science was selected because it is one of the most powerful, up-to-date, comprehensive, and popular scientific literature search engines widely available for analyses of interdisciplinary, peer-reviewed literature [64]. The search terms used in this study were based on those previously used in a number of highly cited literature review papers [65, 66]. The definitions and categorization associated with agriculture and climate change adaptation used by the IPCC [67] were also referenced to establish the inclusion and exclusion criteria, as well as the classifications and sub-categorizations of the literature (Table 1).

**Table 1.**
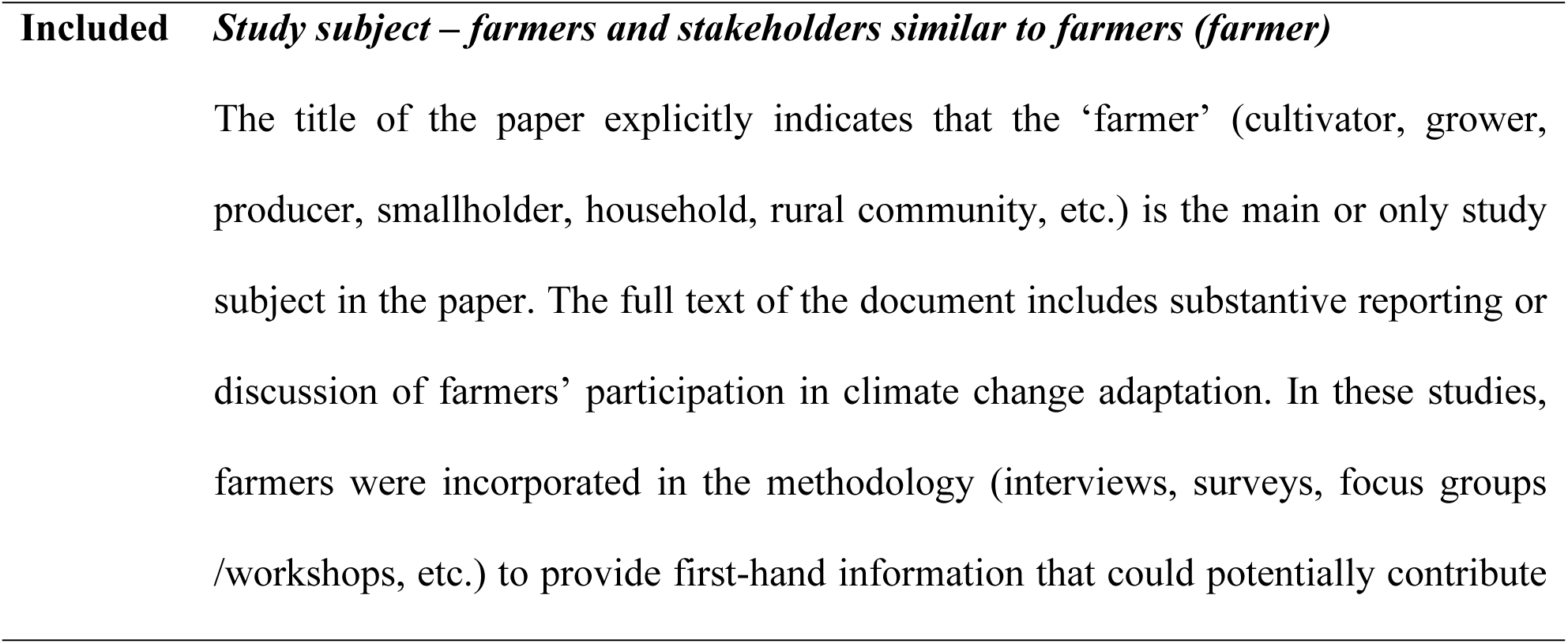
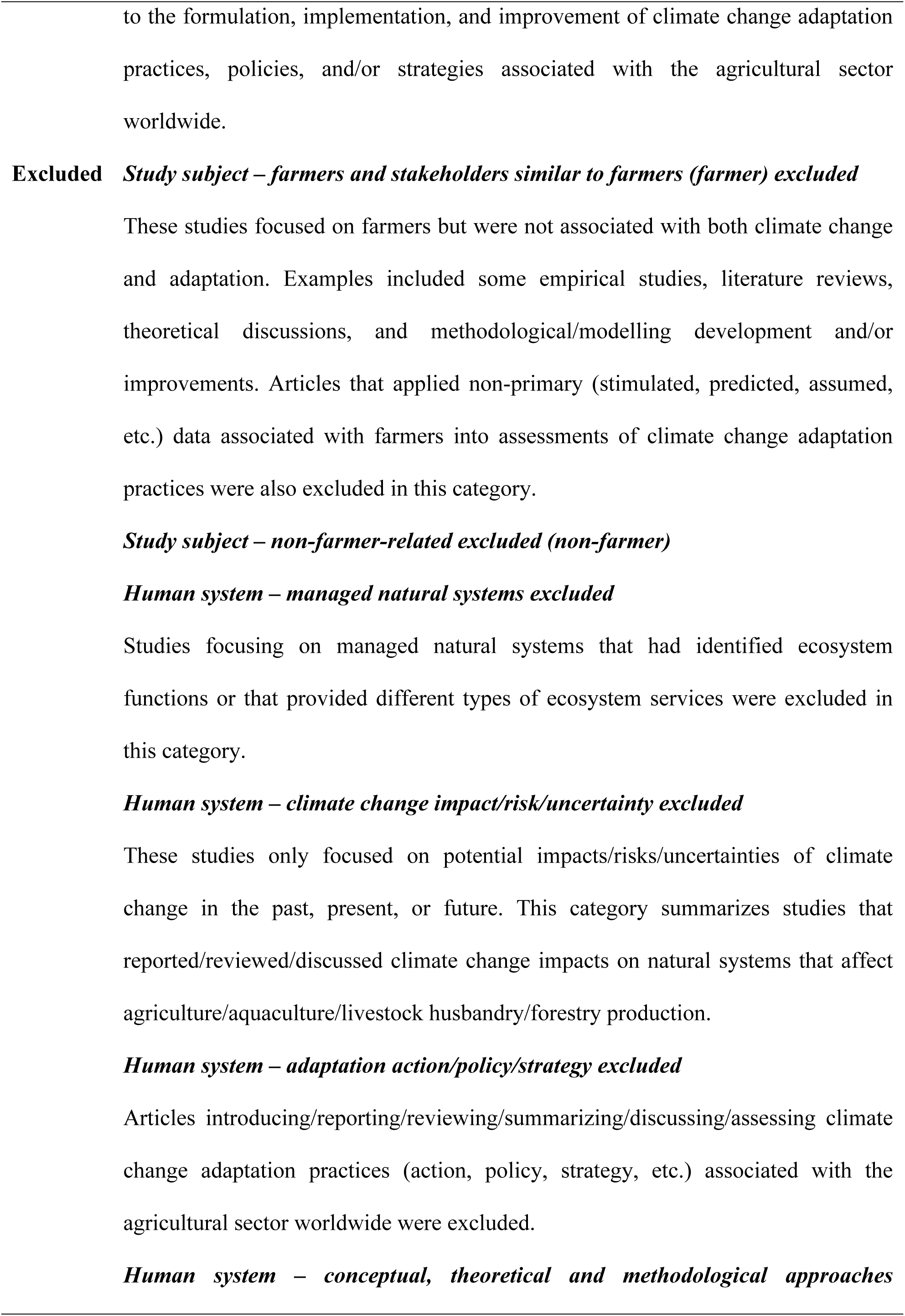
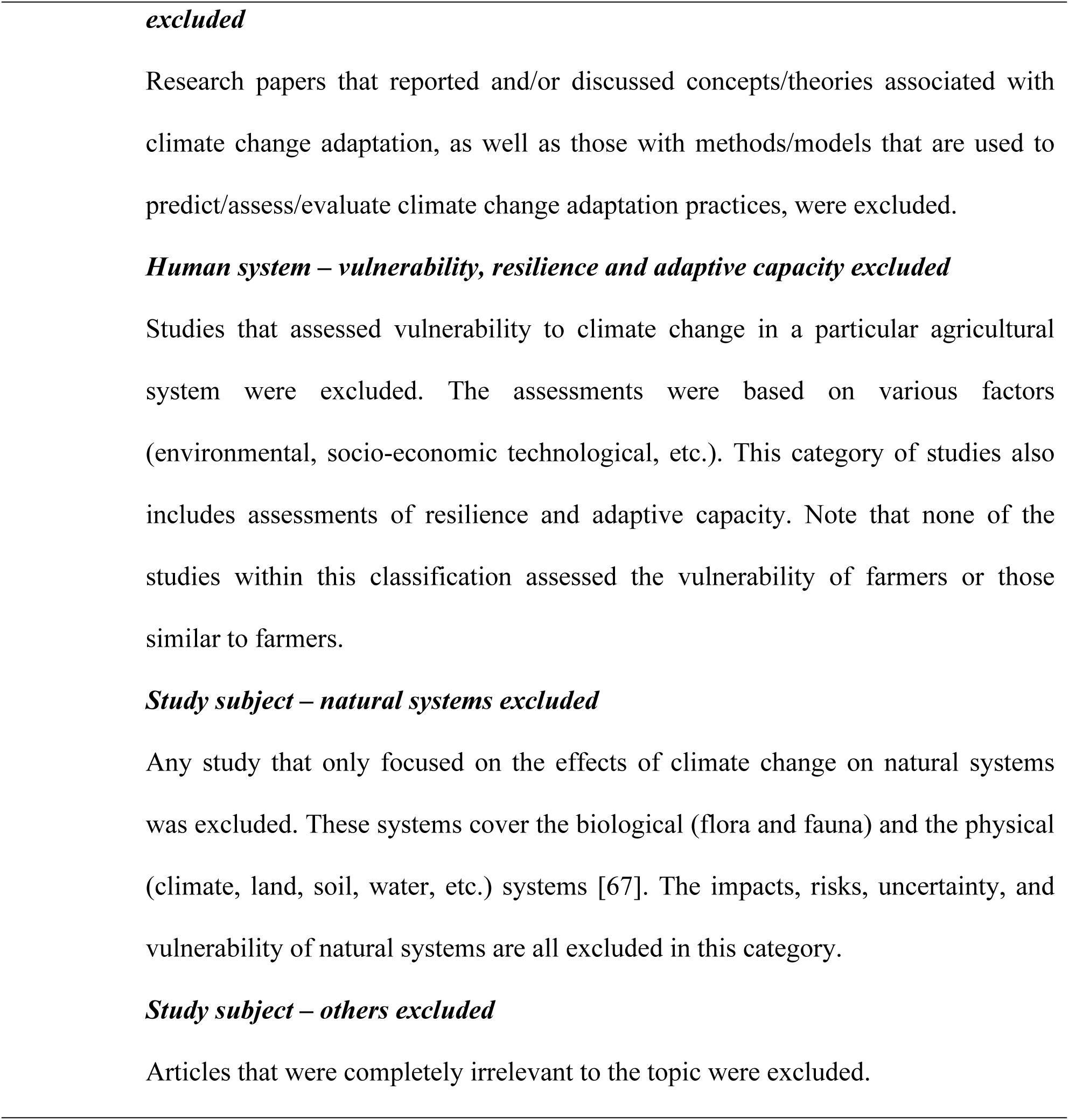
Descriptions of the literature search and inclusion and exclusion criteria classifications.

The literature selection procedure involved three steps. First, we conducted a keyword search in Web of Science. To yield as many relevant articles as possible, this step was conducted in the ***subject*** field. To include variations of the thematic terms, Boolean operators and wildcards were applied to yield the following search terms: “climat* chang*” AND “adapt*” AND “farmer.” We screened for peer-reviewed, English-language literature as a proxy sample of scientific literature. We limited the search to the 2008-2017 period because most of the literature prior to 2008 was already reviewed by the IPCC AR4 (2007), TAR (2001), and other notable review papers [68–71]. Documents in other languages, those outside of the search period or those other than articles and reviews (e.g., abstracts, meeting/conference proceedings, books/book sections) were excluded [65]. This step retrieved 1502 papers.

Second, we conducted a keyword search to further screen the 1502 retrieved papers for relevance in this study. The main purpose was to achieve greater accuracy in identifying only those studies exclusively focused on ‘farmers’ and adaptation to climate change. An effective way to filter out non-farmer-related papers was to conduct a more precise term search in the ***title*** field of papers [33, 66]. To guarantee more precision in the obtained records, we defined the word ‘*farmer*’ as any individual and/or a group of individuals engaged in land-based, cultivation-oriented agricultural production activities that supply basic human needs (e.g., food, fibre), including farmers, cultivators, growers, producers, smallholders, households, and rural communities. These selection criteria narrowed the number of papers to 563.

In the final step, the titles and abstracts of the 563 remaining articles were individually reviewed to evaluate each article’s suitability for inclusion in the final review (Fig 1). We reviewed the ***abstract*** of each paper, specifically looking into the context of research methodology. The aim was to determine whether farmers were incorporated into the methods of each study. Cursory to in-depth full-text reading was performed for those papers without abstracts to evaluate their suitability. Three essential criteria had to be met simultaneously for an article to be included in the final review list: 1) farmers were the only study subject or a primary study subject; 2) farmers were incorporated into the research methodology; and 3) the objectives of studying farmers were closely related to both agriculture and climate change adaptation. Accordingly, 358 papers met all three criteria and were retained for full-text review (Fig 1).

**Figure 1.** Summary of document selection.

### Literature review and analysis

All (n=358) articles and their citations were retained in an EndNote library for external review and validation. Following the methods of Berrang-Ford, Ford (65), a step-wise checklist was developed to survey selected articles, record relevant data, and characterize if and how farmers were studied in the areas of agriculture and climate change adaptation (Table 2). The main purpose for designing this checklist protocol was to standardize article analysis and enable descriptive statistical testing to identify and examine key trends and associations. The findings of the reviewed studies were coded for themes for further analysis.

**Table 2.**
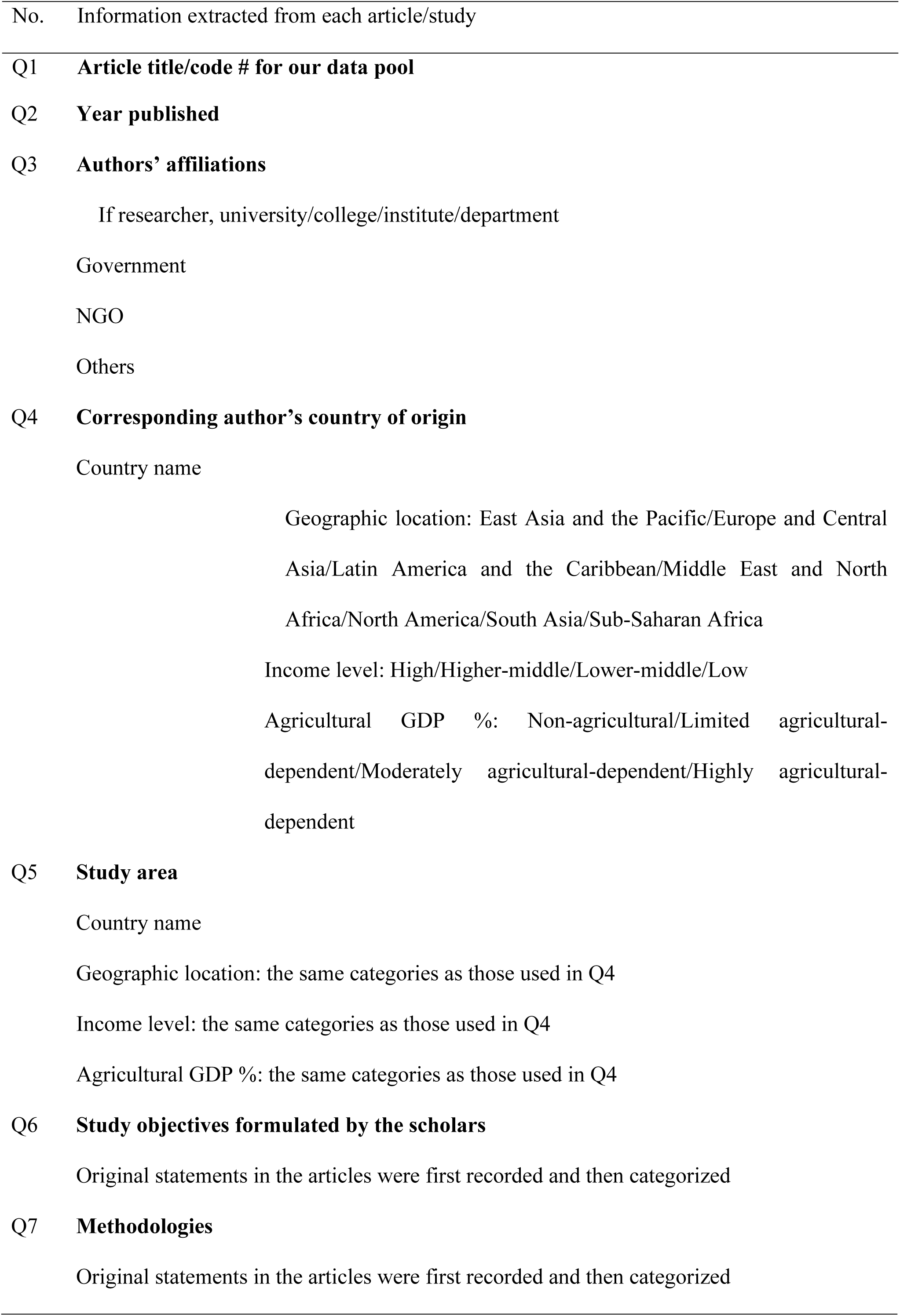
Article review checklist.

|  |  |
| --- | --- |
| No. | Information extracted from each article/study |
| Q1 | Article title/code # for our data pool |

The checklist began with questions about the basic information of each reviewed article, including authorship (e.g., names and affiliations), year of publication, and region of interest. The body of the checklist documented detailed information on each paper, including (1) why researchers were interested in studying/engaging farmers in agriculture and climate change subject areas; (2) the methods used to incorporate farmers in these studies; (3) the countries in which farmers were studied; (4) where the researchers were from; and (5) the researchers’ expertise. The full-text review of the 358 articles was conducted using this checklist.

Descriptive and basic inferential statistics were calculated to identify trends and summarize the quantitative attributes of the data. ArcGIS (ESRI v 10.2) was used for geographic data mapping.

### Secondary data

The geographic distribution of countries was based on the World Bank (WB) classification, in which a total of 217 nations were categorized into seven geographic regions: 1) East Asia and the Pacific (37 nations); 2) Europe and Central Asia (58); 3) Latin America and the Caribbean (42); 4) the Middle East and North Africa (21); 5) North America (3, including Bermuda); 6) South Asia (8); and 7) Sub-Saharan Africa (48). Two indicators were used to categorize socio-economic conditions. The first indicator was income level, based on the 2016 gross national income (GNI) released by the WB. Each country is grouped into one of four categories: low-income, lower-middle-income, higher-middle-income, and high-income. For the present analysis, we considered the countries in the first two categories to be ‘lower-income nations’ and the countries in the last two categories to be ‘higher-income nations.’ The second indicator was the average agricultural-associated GDP (%) of each country within the literature search period (2008-2017). Data were collected from the WB’s online database with agricultural added value (% of GDP) for all but six countries (Table 3). We used these data to group countries with available data into four categories: non-agricultural (<1% GDP from agricultural sector); limited agricultural-dependent (1-10% GDP); moderately agricultural-dependent (11-30% GDP); and highly agricultural-dependent (>31% GDP). For subsequent analysis, we refer to the first two categories as ‘low agricultural-dependence countries’ and the other two as ‘high agricultural-dependence countries.’

**Table 3.**
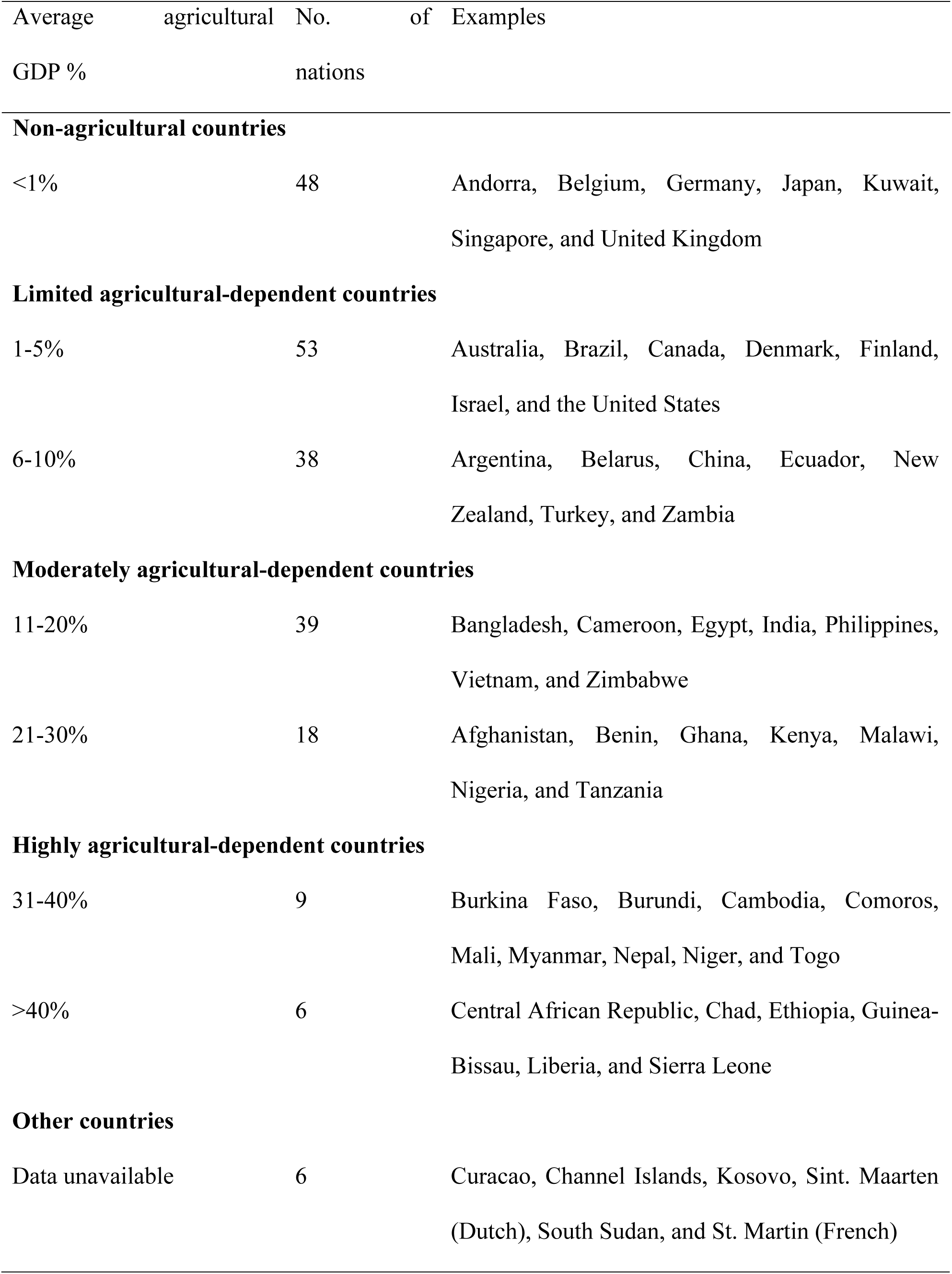
Classifications of countries based on agricultural % GDP.

## Results

### A small but increasing number of studies focus on farmers in climate change adaptation

Of the 1502 initially screened articles addressing farmers and climate change adaptation-associated topics between 2008 and 2017, over 60% (n = 939) focused on the latter, with only peripheral interest in farmers (Figure 1). Of the remaining 563 studies that focused on farmers, 205 were not directly related to either agriculture (e.g., health, anthropology) or climate change adaptation (e.g., trade, insurance). Ultimately, only 358 articles fit all three of our inclusion criteria, accounting for close to a quarter (23.8%) of the 1502 total articles, which is also only a small portion of the total amount of published literature examining either agriculture and/or climate change adaptation more broadly. However, despite the relatively few publications, the number of these studies has been increasing over time. The annual number of articles increased from only one in 2008 to 107 in 2017 (Figure **2**). Considering that the IPCC published its fifth Assessment Report (AR5) in 2014, we also compared the total number of articles published prior to the AR5 (2008-2013, 85 papers) with the total number published afterward (278 papers; Figure **2**).

**Figure 2.** Annual evolution of published papers incorporating farmers in their methods to study agriculture and climate change adaptation (2008–2017)

### Uneven global geographic distribution of the reviewed studies

In total, 358 papers are from 75 countries across all seven geographic regions, accounting for slightly more than one-third (34.6%) of all countries worldwide (217). However, the number of relevant studies from each region differed considerably (Figure **3**). The region with the highest number of studies is sub-Saharan Africa (172 papers from 23 countries, hereafter recorded as 172; 23), followed by East Asia and the Pacific (84; 14), South Asia (69; 6), Latin America and the Caribbean (35; 12), Europe and Central Asia (24; 13), North America (23; 2) and the Middle East and North Africa (6; 5). The geographic disparity in the number of papers was, however, more prominent within regions, with studies often primarily focused on only a few countries within each region. For example, while the number of studies in sub-Saharan Africa accounted for nearly half of all selected papers, of the nearly 50 countries in this region, the papers were dominated by only six nations (Ethiopia, Ghana, Kenya, Nigeria, South Africa, and Tanzania). In East Asia and the Pacific region, almost half of the papers were from China and Australia. Similarly, over 80% of papers from South Asia were from India, Bangladesh, and Nepal. The United States dominated the studies in North America, while studies of Latin American and Caribbean farmers were mostly from Mexico and Guatemala. There was no evidence of dominance by any particular nation in Europe and Central Asia or in the Middle East and North Africa, but many countries in these regions had no representative studies at all.

**Figure 3.** Distribution of papers studying farmers in relation to agriculture and climate change adaptation. Values reflect the number of publications that focus on farmers from each country.

### Majority of studies conducted by higher-income nations

The corresponding authors of the 358 reviewed articles were from 59 countries with different socio-economic circumstances (Figure **4**a), including 33 higher-income nations and 26 lower-income nations. For the numbers of publications in each income category, most papers were from high-income nations (181 papers from 22 high-income nations, hereafter 181; 22), followed by lower-middle-income nations (81; 17), higher-middle-income nations (65; 11), and low-income nations (31; 9). While authors from high-income nations produced over half of the total papers, they were largely concentrated in the United States (53 papers) and Australia (33). For higher-middle-income nations, approximately two-thirds of the papers were written by authors from China (23) and South Africa (18), while over 40% of papers from lower-middle income nations were written by authors from Nigeria (19) and India (15). For papers from authors in low-income nations, more than one-third were from Ethiopia (11). Overall, the top ten nations for corresponding authors were the United States (53), Australia (33), China (23), Germany and Nigeria (19 each), South Africa (18), the United Kingdom (16), India (15), Ethiopia (11) and Japan (10); seven of these countries were higher-income nations.

**Figure 4.** (a) Nation of origin for corresponding authors across different national income levels; (b) Study locations (farmers’ locations) across different levels of national agricultural dependence.

### Farmers from more than half of highly agricultural-dependent countries were not studied

The study locations of the 358 reviewed articles include 75 countries across the full range of agricultural dependence (% GDP, Fig 4b), including 37 countries with low agricultural dependence and 38 with high agricultural dependence. Specifically, the plurality of papers investigated farmers from moderately agricultural-dependent countries (210 papers; 31 countries), followed by farmers from limited agricultural-dependent countries (137; 32), highly agricultural-dependent countries (54; 7), and non-agricultural countries (12; 5). Farmers from the following ten nations were most frequently studied within the review period: Nigeria (25), India (24), China and United States (22 each), Ethiopia and Ghana (19), Australia, Bangladesh and Nepal (16 each), and Kenya (15). Only three of these ten countries are characterised by low agricultural dependence. Although the number of papers focusing on farmers from countries with high agricultural dependence was considerably higher, farmers from nearly half (47%) of countries with high agricultural dependence were not studied at all (34 out of 72). When only the 15 nations that are highly agricultural-dependent (>31% agricultural GDP) were examined, the farmers from more than half (8 nations; 53%) were not studied at all, namely, Burundi, Central African Republic, Chad, Comoros, Guinea-Bissau, Liberia, Sierra Leone, and Togo (see Appendix).

### Diverse objectives and methods of incorporating farmers

We identified five main categories of rationales for why these studies incorporated farmer-focused methods (Figure **5**): 1) understanding farmers’ climate change perceptions (n = 207, 58%), 2) exploration of farmers’ interests in climate change policy and/or decision-making processes (n = 102, 29%), 3) evaluation/examination of farmers’ adaptation performance (n = 100, 28%), 4) assessment/improvement of farmers’ adaptive capacities (n = 90, 25%), and 5) identification of the factors that influence farmers’ choices with respect to adaptation actions (n = 53, 15%). However, the majority of all 358 reviewed studies stated more than one of these five core objectives.

**Figure 5.** Number of papers that formulated different objectives of studying farmers (values are not cumulative)

Notwithstanding the disparity across geographic and socio-economic conditions, social surveys were the most common methodology used to incorporate farmers into these studies. Over two-thirds (n = 243, 68%) of the reviewed studies described this method in their papers, and most of these surveys were conducted in a face-to-face manner (n = 221, 91%), with only 23 of these studies (9%) being conducted remotely. However, all studies that conducted remote social surveys of farmers were from high-income/limited agricultural-dependent nations. These 23 remote social surveys were conducted in three ways: through the mail, on the internet, or over the telephone. Nineteen of these remote surveys (83%) used only one of the three methods, including 9 mail surveys (39%), 6 telephone surveys (26%), and 4 internet surveys (17%). Four surveys (17%) were conducted via both the internet and the mail. The most frequently used survey instrument across all of these studies was a questionnaire (n = 151, 42%), whether following a structured (76, 50%) or unstructured (51, 34%) format; the other 24 (16%) studies did not specify the types of questionnaires that were used. Studies also incorporated farmers by means of other research methods, including key informant interviews (n = 120, 34%), focus group discussions (FGD) (n = 111, 31%), stakeholder workshops (21, 6%), training programmes/experiments (6, 2%), and ethnographic methods (5, 1%).

### Multiple disciplinary interests in studying farmers in agricultural adaptation to climate change

Similar to the literature review performed by Aleixandre-Benavent, Aleixandre-Tudó (33), Berrang-Ford, Ford (65), we also used authorship affiliation to identify those who are interested in studying farmers in agricultural adaptation to climate change. Because this study only focused on peer-reviewed articles, authors are primarily from research organizations (e.g., universities, research institutions). Contributions by authors with non-research affiliations (e.g., government, civil society, NGOs) were minimal due to the study design. In total, only 13 (3.6%) out of 358 reviewed articles involve authors with non-research affiliations. Nevertheless, we found that paper authorship was largely multidisciplinary (Figure **6**), with nearly 60% (n = 212) of studies being the result of collaboration among researchers from different fields. Overall, the academic disciplines (as defined by Science Direct) with the most representation among authors were the agricultural and biological sciences (n = 172, 48%), environmental sciences and ecology (n = 164, 46%), social sciences/economics/humanities (n= 135, 38%), and decision sciences (n = 102, 28%). There were also authors from geography (n = 41, 12%), food science, technology, and policy (n = 30, 8%), and health sciences (n = 8, 2%).

**Figure 6.** Number of papers published by authors with different research disciplines (values are not cumulative) *Others include geography, food science, technology and policy, and health sciences.

## Discussion

### Farmers are the key stakeholders most often overlooked in agricultural climate adaptation research

Climate change has massive impacts on the planet, and both physical and socio-economic systems are severely affected [3, 72]. Among these systems, the agricultural sector is of significant concern [73], and due to population growth and the severe consequences of food scarcity [23], adaptation is necessary and urgent [74, 75]. To adapt to climate change, sustainable farming around the world must necessarily undergo continual adjustments [76]. Because of the necessity and urgency for agriculture to adapt to climate change, both decision-makers and scholars need to recognize the key role that farmers play in the process [77, 78].

Farmers are important stakeholders in agriculture, and their livelihoods are particularly vulnerable to climate change [22]. A comprehensive understanding of farmer-level decision-making processes in various contexts substantially contributes to both the development and implementation of efficient adaptation action plans, strategies, and policies [58, 79, 80]. Previous studies have pinpointed that farmers who have personally observed or who have knowledge of the impacts of natural phenomena and human activity on the environment are likely to be certain about the occurrence of climate change, believe in the dangers posed and are consequently more proactive in accepting new policies and adopting adaptation practices [81, 82].

There are a number of review studies that have focused on agriculture and climate change [41, 83–87], but few have focused specifically on farmers and climate change adaptation [59, 88]. The five published IPCC Assessment Reports have all emphasized the importance of the agricultural sector on climate change adaptation (see, for example, AR3 Working Group II, chapter 5; AR4 Working Group II, chapter 5; AR5 Working Group II, chapter 7), yet they have given minimal to no attention specifically to farmers. In the most recent review by Aleixandre-Benavent, Aleixandre-Tudó (33), the authors found that ‘farmers’ were not among the most frequently selected study subjects in scientific research on climate change in agricultural subject areas.

Moreover, despite the increasing number of studies that have investigated agricultural adaptation to climate change over the last decade, knowledge gaps remain in the existing literature. In particular, the value of the social sciences in agricultural climate adaptation research is recognized but not comprehensively incorporated into these studies [41]. For example, of the studies that have incorporated the farmers’ perspectives, surveys have been used to assess farmers’ perceptions and adaptation practices. However, most of these studies were conducted by natural scientists, and their methods were not as robust as those typically practised in the social sciences, such that the incorporation of farmers in their methodologies is only a tangential part of their study. Consequently, this approach has inherent limitations when researchers seek a broader understanding of farmers’ perspectives and/or behaviours. Thus, more robustly incorporating the methodological approaches of the social sciences as well as those of the traditional natural sciences will help augment the missing and under-represented knowledge and perspectives of farmers in research on agricultural adaptation to climate change. It will also help guide researchers and policy-makers on the global scale. Thus, future research studies should seek to specifically incorporate the perspectives of farmers in more integrated, interdisciplinary policy-making frameworks.

### The existing knowledge on farmers is limited

Despite the methodological imbalance between social and natural sciences, there is an increasing tendency to collaborate across disciplines. For example, the majority of reviewed studies on farmer adaptation to climate change were the product of scholars with multi-disciplinary backgrounds (Section 3.5), and the objectives and methods used to incorporate farmers were diverse (Section 3.4). Nevertheless, the existing studies present clear disparities not only methodologically [41] but also in terms of geographic distribution and socio-economic circumstances [65, 66, 89]. Notably, the geographic distribution of papers is also associated with geographic differences in vulnerability [90]. The number of papers from more vulnerable countries/regions (e.g., Africa, Asia, Latin America) is higher than that from less vulnerable countries/regions (e.g., Europe, North America). Nevertheless, farmers from nearly two-thirds of the countries worldwide have not yet received any attention from the research community, at least within the English-language, peer-reviewed journals that are of most significance to the global scholarly debate [91]. Many of the neglected countries are lower-income nations with high vulnerability to climate change and high rural population ratios. Agriculture plays an out-sized role in these countries’ economies and local livelihoods [92]. This is a major weakness of the existing body of scientific knowledge for a number of reasons.

First, the underrepresentation of farmers from lower-income/highly agricultural-dependent nations in the existing English-language scientific literature limits our ability to comprehensively understand these key stakeholders in a global context. Certainly, such knowledge underrepresentation is partly due to the selection criteria of reviewed articles applied in this study. The scarcity of papers from West Africa, Latin America, Central and East Asia, and Europe is likely because these regions are dominated by non-English-speaking countries (e.g., countries that speak French, Spanish, Japanese, Korea, Chinese). For example, over thirty-three thousand Chinese-language articles were found using the same search criteria via CNKI (the most widely used scientific literature search engine in China). This highlights the ongoing difficulty facing global scientific understanding due to continued language segmentation of the world’s scholarship. Thus, whenever possible, future review studies should be conducted in multiple languages to facilitate improved understanding of current global scholarship.

Second, due to the reliance of agriculture on natural resources and weather, it is inherently vulnerable to the impacts of climate change [21, 93]. However, such vulnerability varies considerably from country to country. For example, rare catastrophic events can cause severe impacts on farmers’ livelihoods in nations across the globe [94], but climate change impacts appear to be most severe on the agricultural sectors and farmers in lower-income/highly agricultural-dependent countries because these countries lack resilient infrastructures [95, 96] and non-agricultural resources [97] and have limited adaptive capacities [98, 99]. Because one of the merits of scientific research is to provide rationales for and stakeholder perspectives that support policy-/decision-making [100–102], how could policy-/decision-makers in these countries achieve these goals without the perspectives of the key stakeholders or even the knowledge of how to reach to them?

Third, analysis of the corresponding author affiliations shows that the current research on farmers and climate change adaptation is driven primarily by scholars in higher-income countries (e.g., the United States, Australia, Germany, the United Kingdom, China). Some of these studies focus on farmers from lower-income/highly agricultural-dependent nations. Nevertheless, the methods to incorporate farmers in studies from higher-income nations are more flexible (such as remote surveys), likely due to the farmers’ higher education levels and access to better infrastructures. Hence, studies of farmers in higher-income nations are capable of yielding larger sample sizes and more easily producing publications. For example, twelve papers were published using the same data from a single mail survey of nearly five thousand farmers from corn belt states in the United States from 2013 to 2017 (see, for example, Arbuckle, Morton (17), Arbuckle, Hobbs (103), Arbuckle, Tyndall (104), Morton, Hobbs (105)). Moreover, higher-income nations are more capable of investing and more likely to invest in research and the subsequent translation of the results into policies [106]. In contrast, lower-income nations have more financial constraints, are less likely to conduct the relevant research and are less able to make and implement research-based policy guidelines [106, 107], but these countries’ economies are more highly agricultural-dependent. Due to the aforementioned education and infrastructure constraints, studies of farmers in lower-income/highly agricultural-dependent countries tend to be conducted with smaller sample sizes, mostly on the village/community scale rather than on the national scale [108–110]. Although there are several highly cited papers from lower-income countries [111–113], the existing scholarship concerning farmers from these countries is considerably smaller than that from higher-income countries.

Because farmers from nations characterised by high agricultural dependence have been underrepresented over the last decade in English-language studies of agricultural adaptation to climate change, we propose that future research collaborations should specifically seek to investigate farmers from these nations to help fill this void. Fortunately, although the overall representation of studies from highly agriculturally dependent countries is relatively low, we found that a few nations (e.g., Nigeria, India, Ethiopia, Ghana, Bangladesh, and Nepal) have greater representation than do most less agriculturally dependent countries. Nevertheless, farmers from eight countries in the most critical, highly agriculturally dependent countries (>31% agricultural GDP) were not studied at all: Burundi, Central African Republic, Chad, Comoros, Guinea-Bissau, Liberia, Sierra Leone, and Togo. Special efforts should be made to incorporate the farmers from these nations into future studies.

### Challenges and opportunities for future studies

In 1997, for the first time in history, the international community reached an agreement to address climate change, namely, the Kyoto Protocol (KP), which clearly articulated that the impacts of climate change can be addressed through mitigation efforts. Although there have been many debates among scholars and governments about the KP, it helped to determine that a reduction in GHGs should be the primary objective in humans’ response to climate change [114]. However, by 2013, when the U.S. National Oceanic and Atmospheric Administration (NOAA) documented the surpassing of 400 ppm of carbon dioxide in the Earth’s atmosphere, two clear messages were gleaned: first, mitigation through GHG reduction has limits and may have already failed to cope with the changing climate, and second, the impacts of climate change on human development will likely continue and become even more severe in the future. Consequently, research and political agendas have begun to evolve to prioritize adaptation efforts [115, 116], especially in key sectors, such as agriculture, and among key stakeholders, such as farmers.

Based on our findings, the key role that farmers play in agricultural adaptation to climate change has begun to be recognized globally [117–119]. An increasing number of research collaborations have been conducted across various academic disciplines [120, 121], geographic locations [122, 123], and socio-economic circumstances over the last decade, but these collaborative efforts must continue. The current explorations and understandings of farmers’ roles in agricultural adaptation remain limited. A significant number of farmer-related studies are not actually focused on farmers [124–126]. For example, in some crop modelling analyses, farmer attributions are applied as input data, but many of these data are only hypothetical assumptions [127]. Whether farmers are aware of and/or engaged in research to share their perceptions, opinions, and other attitudes towards climate change adaptation remains unclear in most otherwise relevant studies. Consequently, existing knowledge of farmers can hardly equip policy-makers and/or scholars to reach the objective of ‘establishing a global goal on adaptation’ (Article 7.1, the Paris Agreement, 2015) for the agricultural sector. However, current efforts to incorporate farmers in climate change adaptation studies have already laid the foundation for future scholarship from more diverse backgrounds and multiple languages to fill the knowledge gaps and expand the research context more broadly.

## Conclusions

Despite the modest but growing number of research studies focusing on farmers in agricultural adaptation to climate change over the last decade, particularly since 2015, we found several key weaknesses in the current English-language published literature. First, the number of studies that specifically focus on farmers in climate change adaptation is still relatively small compared to the total number of otherwise relevant studies; second, the current understanding of farmers’ perspectives on climate change adaptation is inequitable on the global scale. Third, most existing studies have been conducted by researchers from higher-income nations. However, these researchers have focused more on farmers in lower-income nations than on farmers from their own countries, likely because farmers in lower-income nations are more vulnerable to the impacts of climate change; thus, farmers in many lower-income/highly agriculturally dependent nations have not yet been studied in English-language scholarship. Therefore, future studies should particularly seek to focus on farmers from these countries or to otherwise publish their findings in multiple languages. Due to ongoing challenges associated with the segmentation of scholarship by language, future reviews should also be conducted in multiple languages to broaden global understanding. Moreover, the subjects of agriculture and climate change adaptation are usually categorized within the natural sciences, yet our review indicates that the methodologies of the social sciences appear to provide the most efficient tools to incorporate one of the most important stakeholders for both agriculture and adaptation to climate change: farmers. Therefore, greater collaboration among scholars from the natural and social sciences should be promoted in the future.

## Acknowledgements

The authors wish to thank all who offered comments and helpful feedback during the numerous brainstorm sessions and group discussions. The authors thank Professor Patrick Regan of University of Notre Dame for sharing his constructive and meaningful ideas.

---

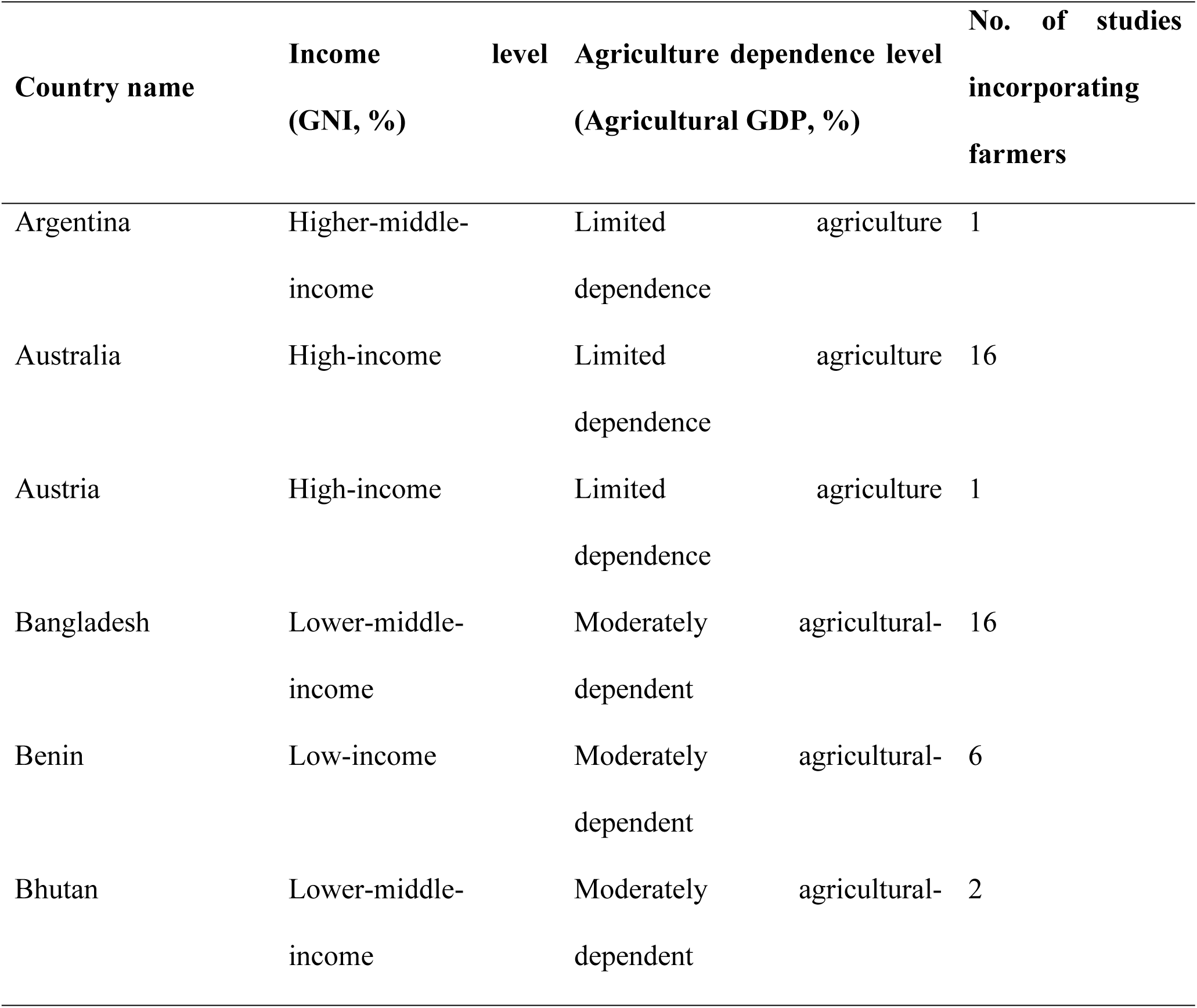
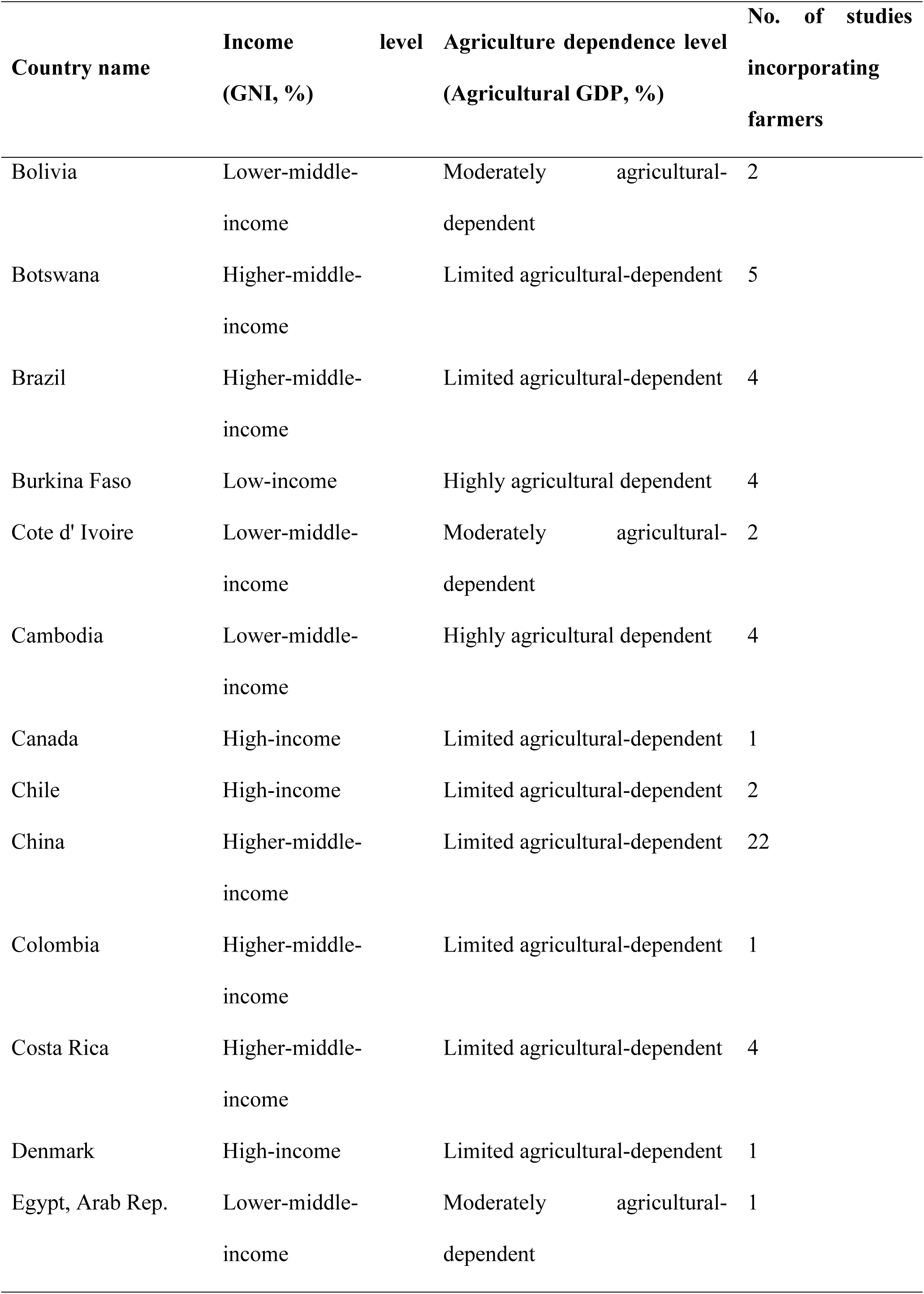
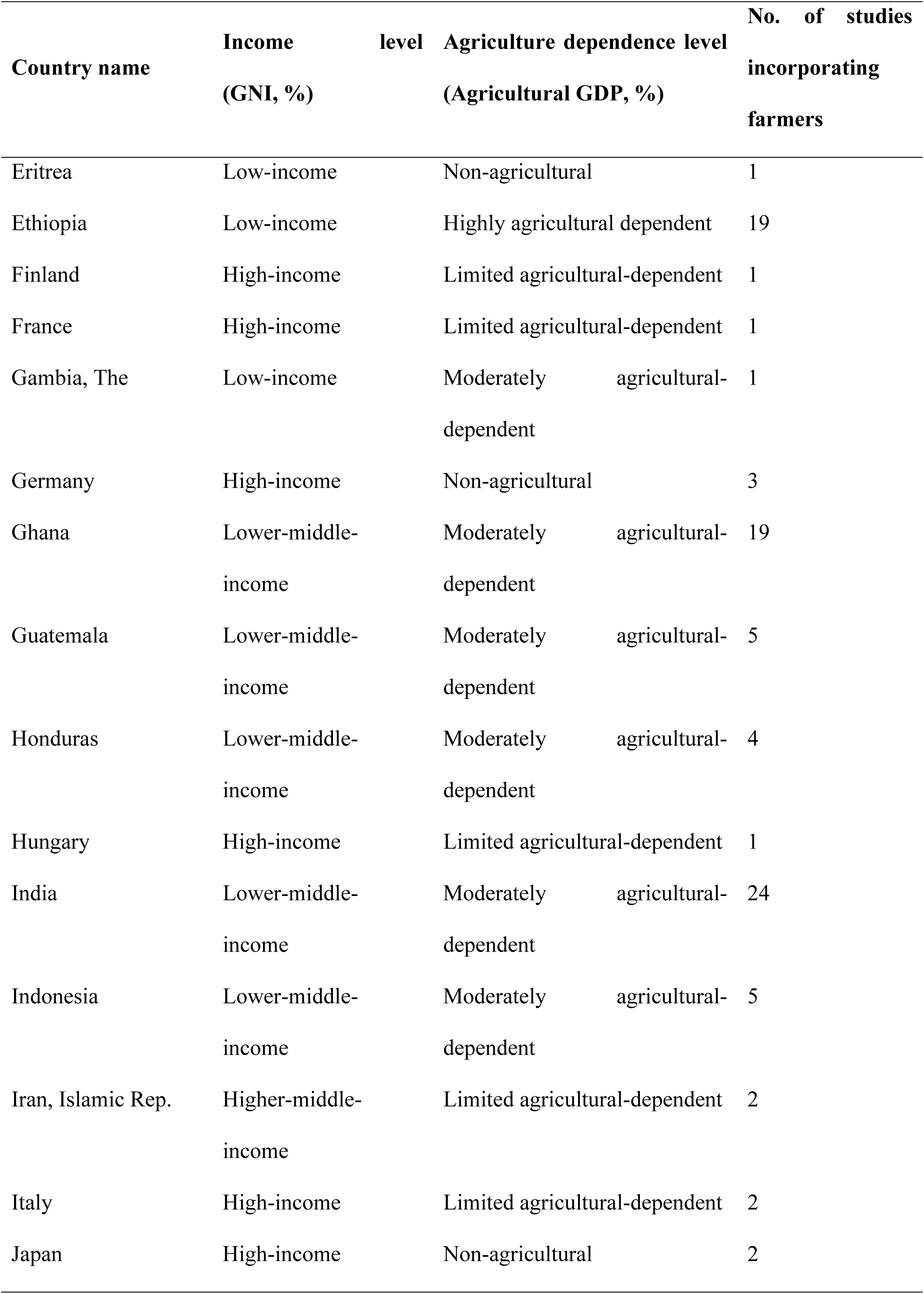
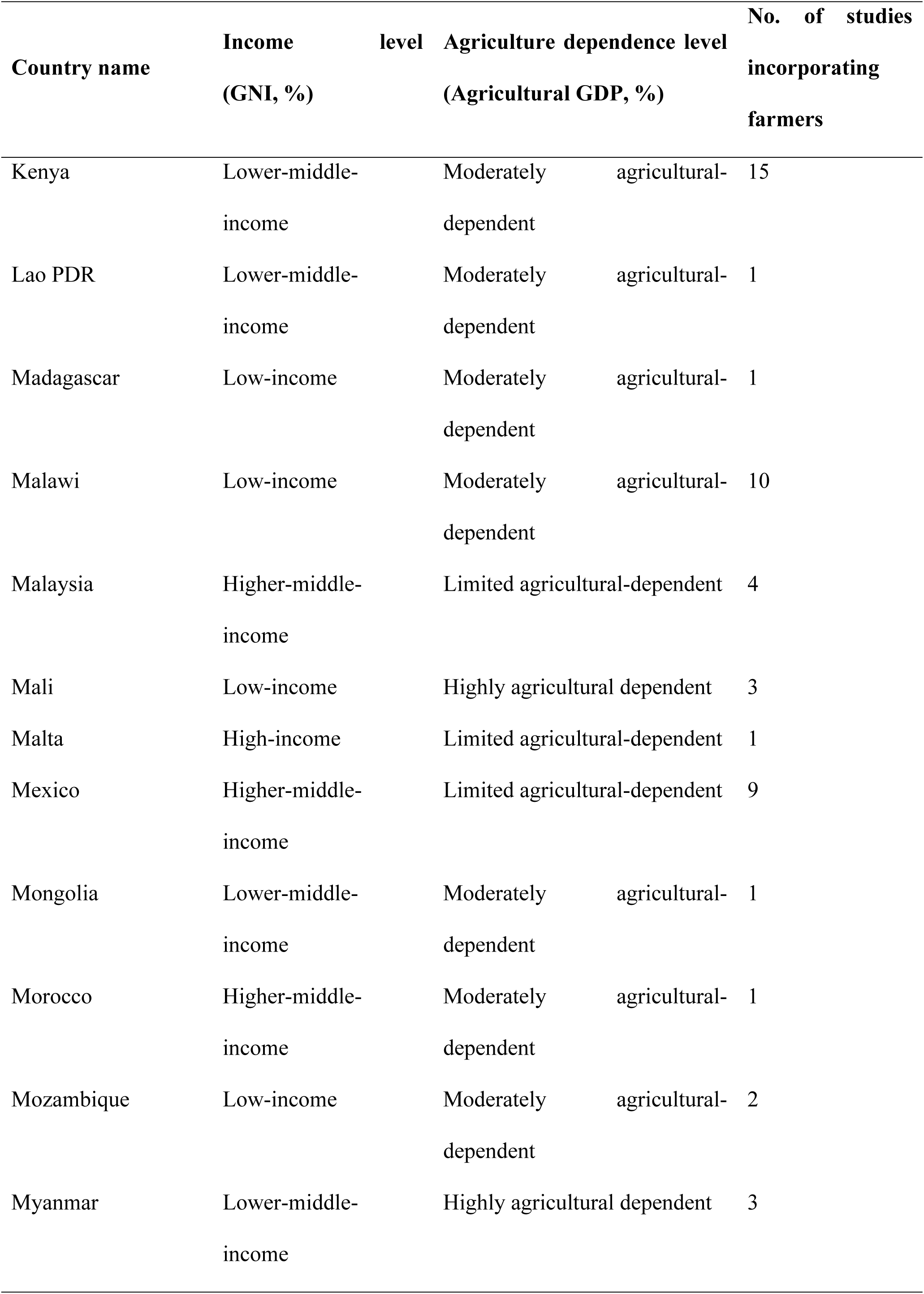
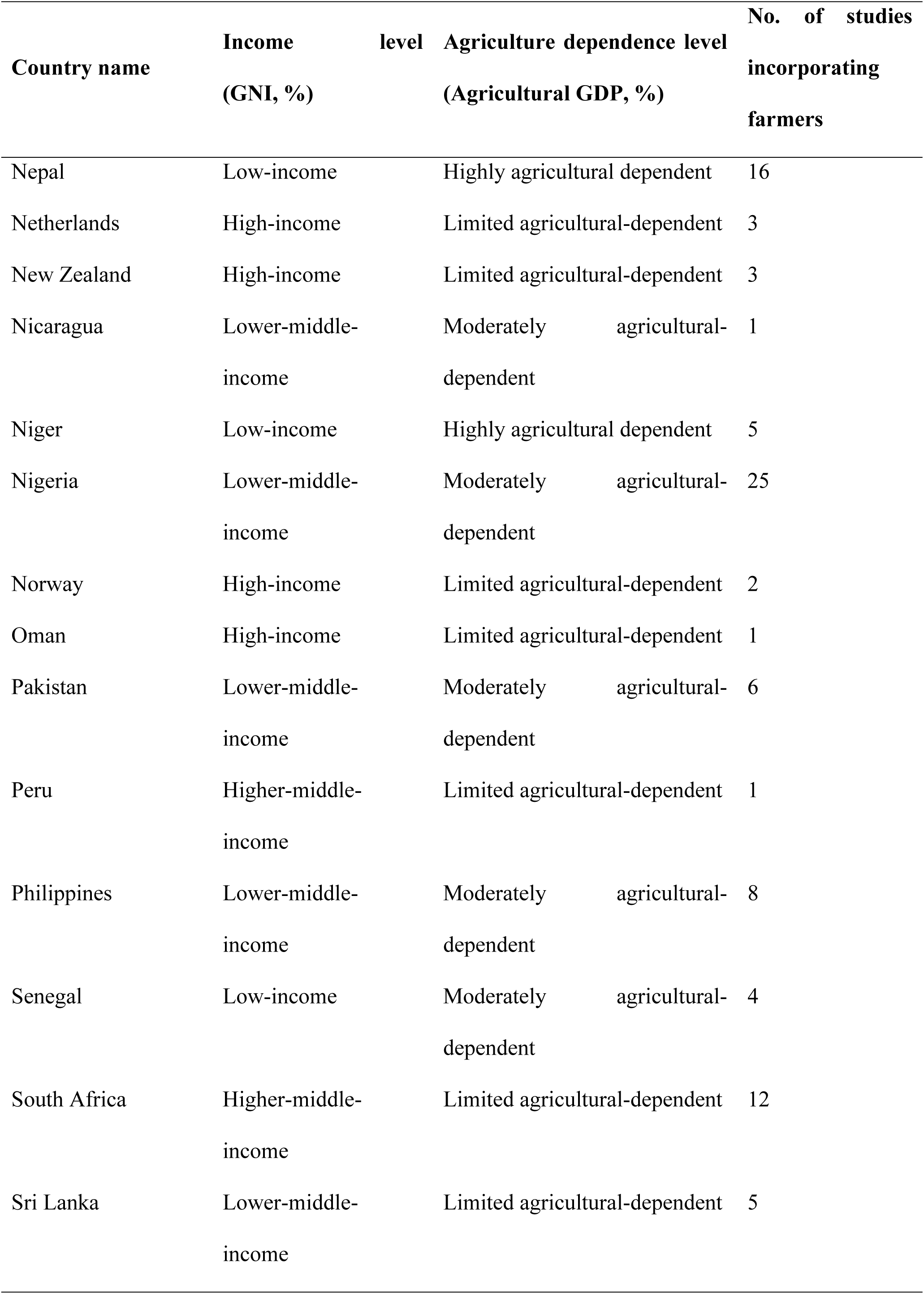
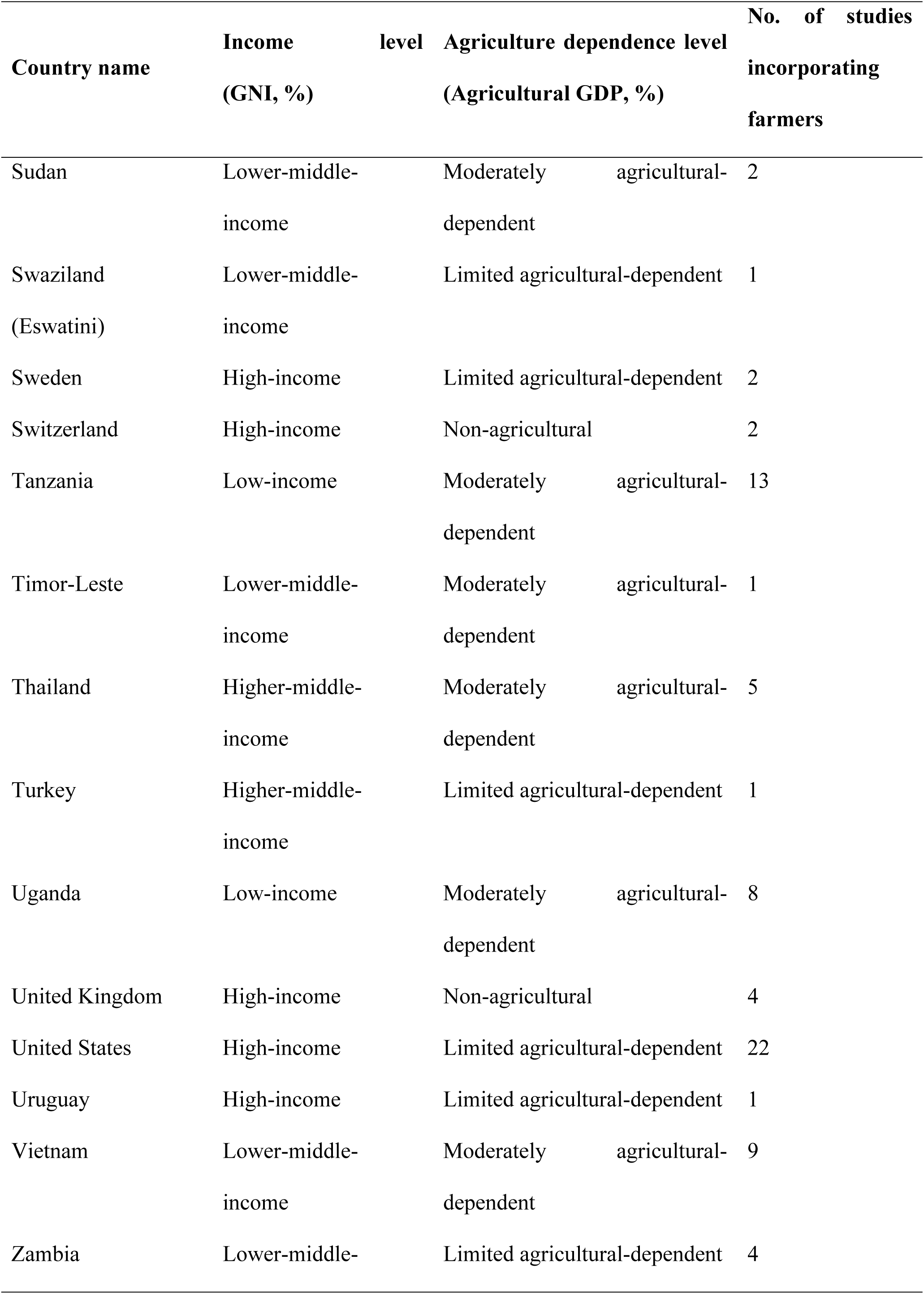
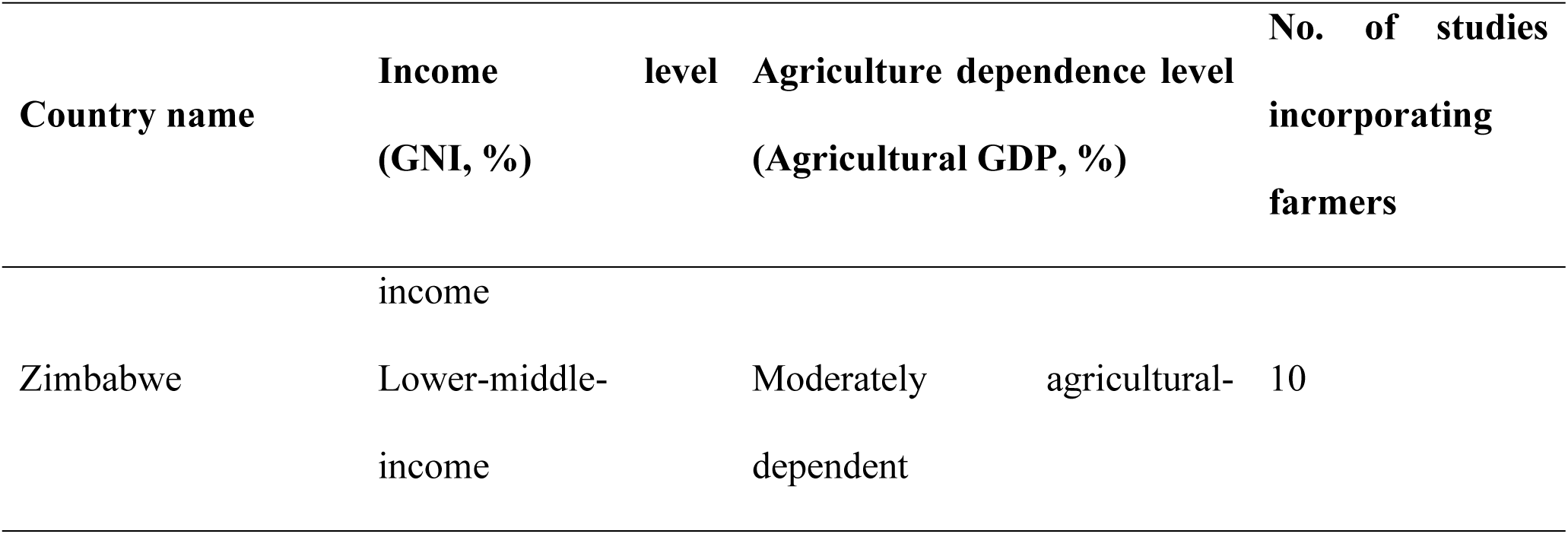
Appendix: Number of studies incorporating farmers from each of the 75 countries.

